# Dynamic Magnetic Resonance Imaging of Whole-Stomach Motility in Rats

**DOI:** 10.1101/2025.06.03.657757

**Authors:** Xiaokai Wang, Ulrich M. Scheven, Zhongming Liu

## Abstract

**Objective:** Understanding gastric physiology in rodents is critical for advancing preclinical neurogastroenterology research. However, existing techniques are often invasive, terminal, or limited in resolution. This study aims to develop a non-invasive, standardized MRI protocol capable of capturing whole-stomach dynamics in anesthetized rats with high spatiotemporal resolution.

**Methods:** Experiments were performed in a 7-Tesla MRI system. Gadolinium-doped test meals were prepared to enhance intraluminal contrast in T_1_-weighted MRI. Based on a modified multi-slice gradient-echo sequence, our protocol integrates respiratory gating to minimize motion artifacts, spatial saturation to improve intraluminal contrast, and slice grouping to optimize the trade-offs between signal-to-noise ratio and motion sensitivity. Image acquisition was accelerated using a time-interleaved k-space undersampling scheme, with missing data reconstructed through k-t interpolation. Image quality and gastric motility were quantitatively assessed.

**Results:** The protocol enabled successful imaging of the stomach and visualization of its pseudo-periodic dynamics in anesthetized rats. The gadolinium-doped meal produced relatively homogeneous intraluminal contrast, allowing clear delineation of gastric anatomy, volume, and motility. The retrospectively reconstructed image exhibited high image quality and yielded reliable estimates of antral contractions, confirming the effectiveness and robustness of k-t interpolation method. Estimated antral contraction amplitude and velocity showed minimal deviations from the reference values, whereas contraction frequency estimation remained highly consistent and accurate. Prospective acquisitions using the accelerated imaging protocol successfully imaged the entire stomach and major intestinal regions, acquiring 24 slices every < 3 s and capturing antral contraction at ∼5 cycles per minute.

**Conclusion:** We established an accessible and standardized imaging protocol that encompasses contrast meal preparation, animal handling and training, and a contrast-enhanced dynamic GI MRI acquisition and reconstruction framework.

**Significance:** This protocol provides a comprehensive, robust, non-invasive tool for studying gastric motility and dysmotility in rodents, offering strong potential to advance preclinical gastrointestinal motility research.

**Graphic Abstract:** In this paper, we report a standardized, non-invasive imaging protocol that encompasses animal handling and training, contrast meal preparation, and a contrast-enhanced dynamic GI MRI acquisition and reconstruction framework for imaging whole-stomach dynamics in anesthetized rats.

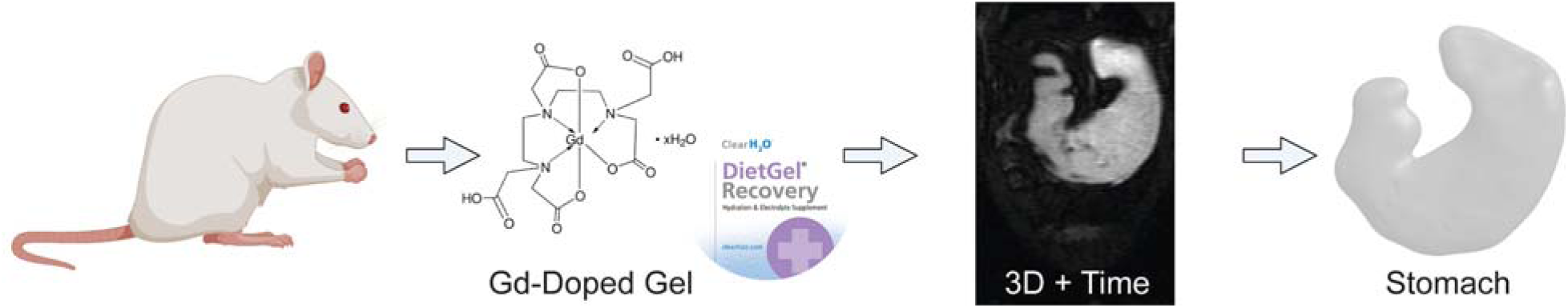

## Introduction

As the gatekeeper for the gastrointestinal (GI) tract, the stomach manages complex motor functions crucial for food intake and digestion (1). Its proper functioning relies on coordinated movements of its regions: fundus, corpus, antrum, and pylorus (2). Dysfunction in gastric motility can lead to significant disorders, such as functional dyspepsia and gastroparesis (3,4). To prevent and manage such disorders, it is critical to characterize gastric physiology and pathophysiology using techniques that can be translated from preclinical to clinical studies.

Rodents, particularly rats, are widely used in animal studies of GI motility due to their anatomical and physiological similarities to humans (5). However, current methods for assessing gastric functions in rats differ significantly from those used in clinical practice, creating translational barriers. Traditional preclinical methods, such as marker-based assays of gastric emptying or food transit (6,7), are terminal, precluding repeated measurements. Alternative methods involving implanted electrodes or strain gauges (8,9) are invasive, potentially disrupting physiology or pathophysiology and confounding experimental outcomes.

MRI has emerged as a promising non-invasive technique for capturing a range of GI functions in both humans and animals (10–14). In humans, MRI has been utilized to evaluate gastric accommodation (15–17), peristalsis (18–21), secretion (22,23), intestinal motility (24,25), and gastric emptying (18,21). Relative to human studies, GI MRI applications in preclinical settings remain limited (11,12,26,27). Previous approaches suffered from either incomplete spatial coverage or compromised speed or resolution, restricting dynamic whole-stomach imaging of gastric motor function.

Advances in accelerated MRI, utilizing parallel imaging (28–30), model-based reconstructions (31–35), and deep learning (36,37), have the potential to overcome existing constraints on spatial coverage and spatiotemporal resolution for dynamic GI MRI. These advancements have not yet been thoroughly applied to preclinical gastric MRI, to our knowledge. Herein, we describe a comprehensive dynamic GI MRI protocol specifically designed to capture gastric motility in rats. Our approach leverages k-t linear predictability to accelerate data acquisition, enabling unprecedented visualization of gastric motility at submillimeter spatial resolution and high temporal fidelity with a whole-stomach field of view (FOV). Our MRI acquisition and reconstruction method, combined with optimized protocols for animal handling and contrast-labeled meal preparation, establishes a standardized, non-invasive imaging platform ideal for preclinical studies of GI motility in health and disease.

## Methods and Materials

### Animals

Fourteen Sprague-Dawley rats (male, 230-430 g, Envigo, IN, USA) were used for this study. All experimental procedures were approved by the Unit for Laboratory Animal Medicine and the Institutional Animal Care & Use Committee at the University of Michigan. Rats were housed under controlled conditions (temperature: 68–79°F, relative humidity: 30–70%) and a 12:12 hours dark-light cycle.

Rats were provided with a standardized test meal, either ingested voluntarily (n = 12) (38) or administered through intragastric gavage (n = 2), 10-30 minutes prior to MR imaging. To facilitate the voluntary ingestion, twelve rats underwent a seven-day diet training protocol before the imaging experiment (11,39). Briefly, during the first two days, rats had ad libitum access to both regular chow and diet gel (DietGel^®^ Recovery, ClearH2O, ME, USA) until 6 p.m. on the second day. From day three to day seven, rats only had ad libitum access to the diet gel from 12 p.m. to 6 p.m., with no food provided outside these hours. Water remained available at all times. Following this training, rats were able to consume ∼5 g of the test meal within 30 minutes after being fasted overnight for 18 hours. For intragastric administration, rats were acclimated to gentle but firm handling and restraining for at least two days before the imaging experiment. They were given a total of ∼3.4 g of test meal through intragastric gavage in up to three doses within 10 minutes.

### Contrast Agent and Test Meal

The test meal was a homogeneous and semi-solid mixture of diet gel and Gadolinium (Gd)-DTPA (#381667, Sigma Aldrich, MO, USA). Specifically, the gel provided 112.4 kcal per 100 g, including 71-75% water, 23.1% carbohydrates, 1.9% fat, and 0.6% protein. We made a 1 mL, 182 mM Gd-water solution with a probe sonicator (Q700, Qsonica, CT, USA), and further mixed it with 25 mL of heated, liquified gel to prepare a 7 mM Gd-doped mixture. The mixture was vortexed and cooled overnight to yield a uniform, solidified test meal (Fig. 1) with a short T_1_ relaxation time (∼17 ms at 7T) (Fig. 2).

**Figure 1.**
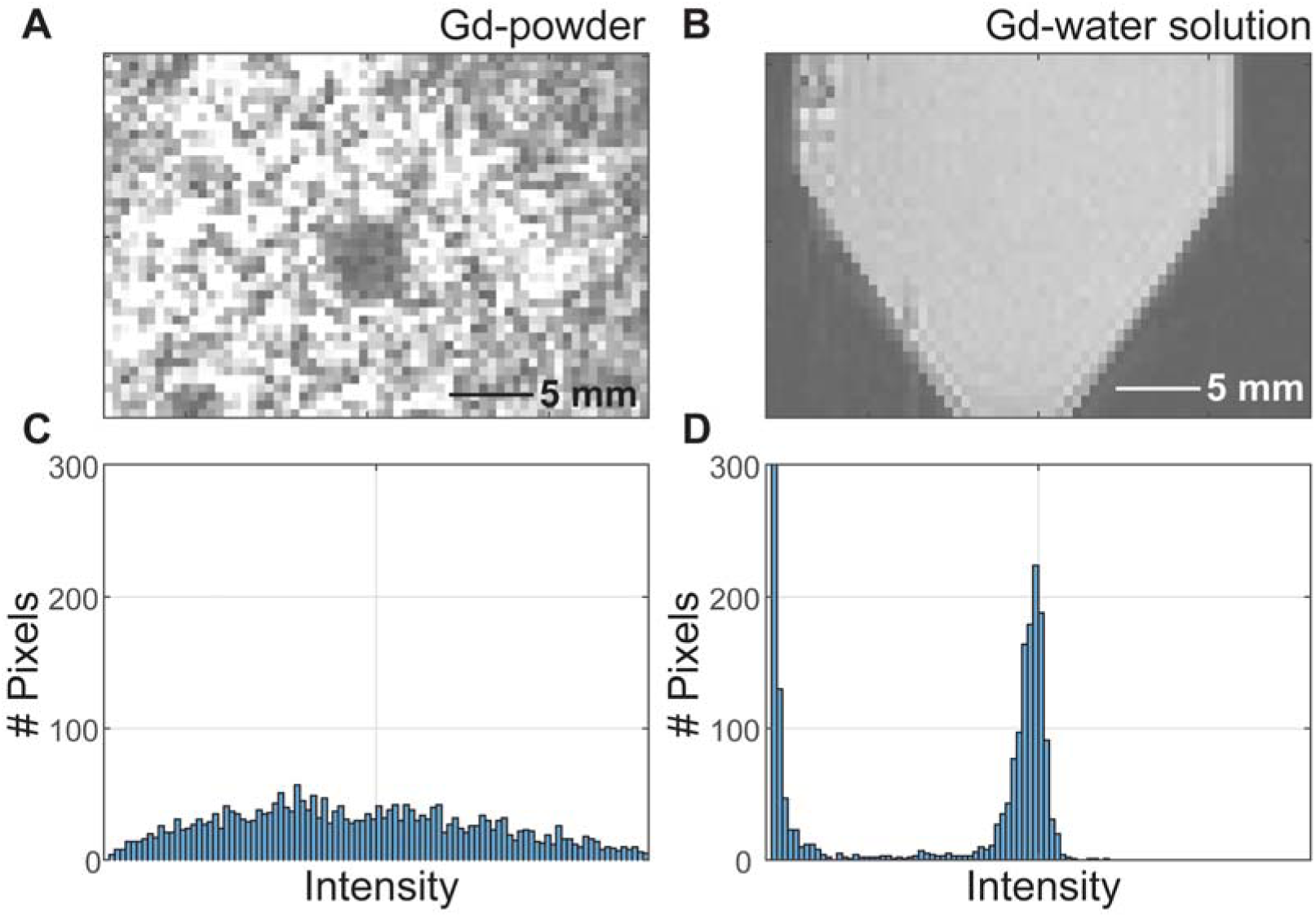
Characterization of Gd-doped meals. **(A)** Diet gel is mixed with Gd-DTPA in powder form (Gd-powder). **(B)** Diet gel is mixed with Gd-DTPA dissolved in water (Gd-water solution). Correspondingly, the pixel intensity distribution of images with Gd-powder and Gd-water solution is in **(C)** and **(D)**.

**Figure 2.**
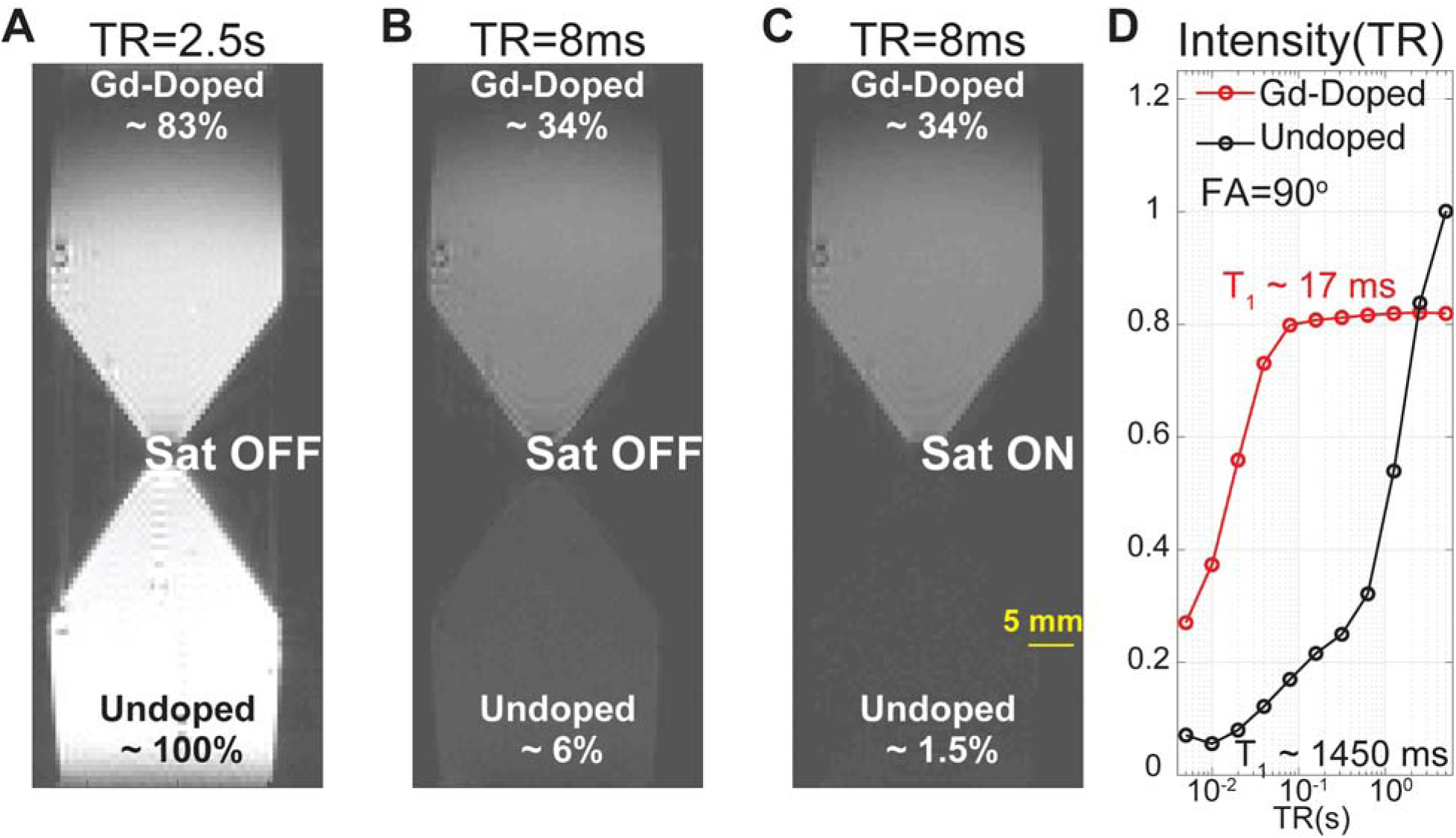
Gradient echo images of undoped and Gd-doped test meals, for different recovery times TR, with and without spatial saturation. **(A)** TR = 2.5s, **(B)** TR = 8ms, **(C)** TR = 8ms with spatially non-selective saturation. **(D)** Pixel intensity of undoped and Gd-doped meals, as a function of TR. Intensities are all normalized to the signal of the undoped meals at long TR.

### MRI Acquisition

Imaging was performed in a 7-Tesla horizontal-bore small-animal MRI system (7T/310/ASR, Agilent Technologies, CA, USA) equipped with a gradient set (inner diameter: 120 mm; strength: 400 mT/m) and a volume transmit and receive ^1^H RF coil (inner diameter: 60 mm). Rats were anesthetized (39) and maintained stable physiology (respiration: 30–60 cycles per min (cpm), heart rate: 220–310 beats per min, body temperature: ∼37°C), monitored via an MRI-compatible system (Model 1030, SA II, NY, USA). The respiratory signal was recorded with a pressure-sensitive pillow placed under the animals. Fig. 3 depicts schematically a typical respiration trace of rats under anesthesia. The image acquisition was triggered by the respiratory signal and confined to the quiescent phase (slow exhalation) to minimize motion artifacts. With a respiration rate of 30-60 cpm, the time window for each MRI acquisition is 1-2 s per respiration cycle.

**Figure 3.**
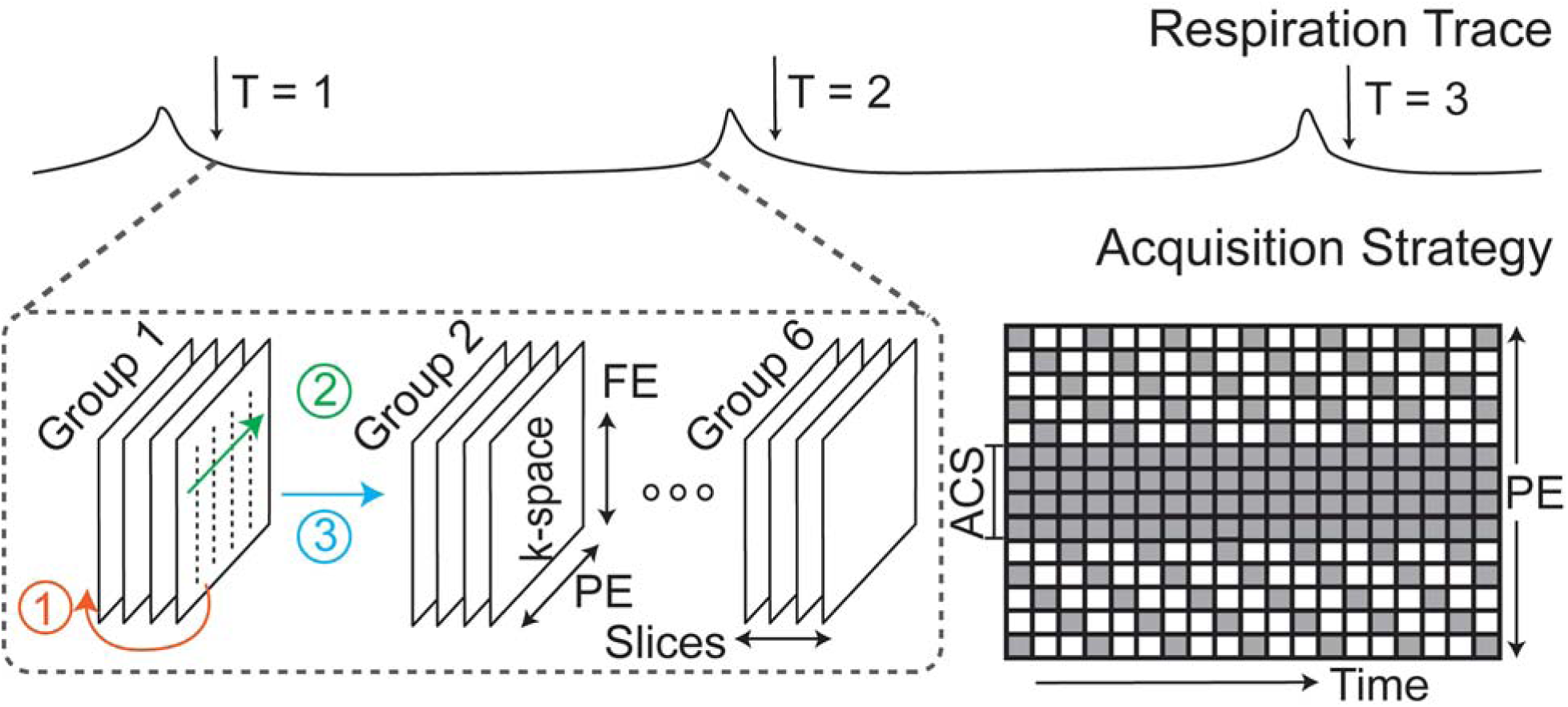
Schematic acquisition strategy. The respiratory signal of anesthetized rats was used to trigger the image acquisition. Each respiration cycle is characterized by a short spike in pressure, due to rapid respiratory motion at that time (inhalation), followed by a longer and reasonably quiescent interval (slow exhalation). Arrows at T=1, T=2, and T=3 mark the gating signal at the beginning of a complete stomach image acquisition. Following the gating signal, 24 slices are acquired in 6 consecutive groups (packets) of 4 adjacent slices, as described in the text. Colored numbers indicate the order of the loop structure, from slice dimension, phase encoding (PE) direction, and groups. The k-space sampling pattern is plotted along PE (vertical axis) and time (horizontal) dimensions. The sparse k-space sampling pattern repeats every three acquisitions in this schematic diagram. Each box is one line of frequency encoding (FE). Gray box: acquired PEs, white box: skipped PEs, ACS: autocalibration signals.

Images were acquired using a hybrid multi-slice gradient-echo sequence with 24 slices (1.5 mm thickness) grouped into six consecutive packets of four adjacent slices each (Fig. 3). This scheme was chosen to ensure signal intensity by using an adequate TR for magnetization recovery, while also minimizing motion artifacts by limiting the time window during which each slice was acquired. Each respiration-triggered acquisition scanned slices sequentially from the ventral to the dorsal side. Acquisition parameters included TR = 10.6 ms, TE = 1.6 ms, flip angle (FA) = 25°, FOV = 64 × 42 mm², matrix size = 128 × 84, and nominal in-plane resolution = 0.5 × 0.5 mm².

Spatially non-selective saturation pulses were applied immediately before each packet of slices to suppress signals from surrounding tissues and flowing blood. This approach enhanced gastric contrast by leveraging the rapid T_1_ recovery of Gd-doped contents (T_1_ ∼17 ms) compared to the much slower recovery of surrounding tissues (T_1_ >800 ms), while reducing motion-related and aliasing artifacts. Given our effective recovery time of ∼127 ms (defined as the interval between the saturation pulse and the gradient echo at the center of k-space), the longitudinal magnetization of gastric content fully recovered, while signals from surrounding tissues recovered only marginally (< 20%) (Fig. 2).

Data acquisition was accelerated using a time-interleaved undersampling scheme in k-space. As illustrated in Fig. 3, phase-encoding (PE) lines (bandwidth: 100 kHz) were partially sampled at consecutive time points, alternating skipped lines to achieve a reduction factor (R) of 3 at the peripheral k-space (12 lines), while auto-calibration signal (ACS, 12 lines) were fully sampled at the k-space center. This sampling scheme balanced the spatial and temporal resolution, enabling full stomach coverage within less than 3 seconds per image volume with respiratory gating.

### MRI Reconstruction

Reconstruction leveraged linear predictability (40) in k-t space to interpolate the undersampled k-space data (Fig. 4), adapting principles similar to k-t GRAPPA (41) for a single-coil scenario. Our justification for linear predictability relied on two key assumptions. First, the spatially uniform Gd-doped gastric contents created regions of piecewise homogeneity in the image space, effectively acting as spatial multipliers analogous to coil sensitivity profiles in parallel imaging, thereby corresponding to convolution kernels in k-space. Second, gastric motility exhibited quasi-periodic motion (∼5 cpm), characterized by narrow-band frequency components, thus justifying the assumption of time-invariant convolution kernels for temporal interpolation. With these considerations, k-t convolutional interpolation kernels derived from fully sampled ACS (12 central k-space lines) could be used to accurately estimate the missing peripheral samples despite an undersampling factor of R=3 or higher. Specifically, we estimated the kernel weights by fitting with ACS and minimizing the least square errors. This approach facilitated robust reconstruction, yielding high-resolution dynamic images with clear delineation of gastric structures and motility.

**Figure 4.**
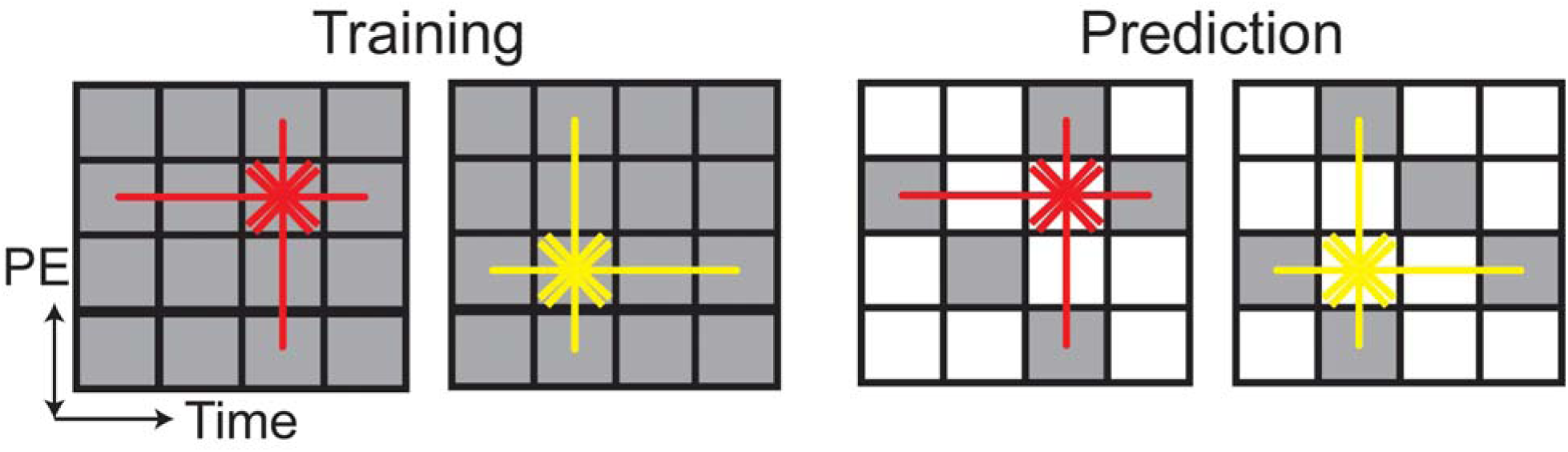
Interpolation kernels for image reconstruction. Take the reduction factor, R = 3, as an example. Two kernels (red and yellow) with distinct shapes were used. During training, acquired PEs at the central k-space (ACS, gray) were used to train the kernel weights to predict the other, following the colored arrows. During prediction, skipped PEs (white) at the peripheral k-space were estimated using the trained kernels, following the colored arrows.

### Validation and Evaluation

Validation and evaluation of the proposed imaging approach were performed through both retrospective simulations and prospective data acquisitions. Retrospective simulations involved applying various reduction factors (R = 2, 3, 4 through 11) and numbers of auto-calibration signal lines (ACS = 12 and 24) to fully sampled reference datasets (four slices with spatial saturation, TR = 10 ms, TE = 1.4 ms, FA = 25°, acquisition time = 0.9 s per image volume).

Reconstruction performance was evaluated by computing quantitative metrics relative to the fully sampled reference images, including relative error (RE) (42), structural similarity index (SSIM) (43), and peak signal-to-noise ratio (PSNR). Before calculating these metrics, images were normalized to [0, 1]. In addition, gastric motility metrics, including contraction amplitude, frequency, and propagation velocity, were extracted from the reconstructed images and compared against fully sampled reference data using the procedures described elsewhere (11). Paired t-tests were performed to evaluate the statistical significance of the differences in motility measures (a significance level of alpha = 0.05).

Prospective acquisitions utilized the proposed accelerated imaging protocol. Reconstructed images were assessed for their ability to accurately depict gastric dynamics and anatomy. Gastric motility metrics were evaluated and compared with the normal range reported elsewhere (11). Gastric volumes measured from time-averaged dynamic images were quantitatively compared with anatomical reference scans (same slice thickness, no spatial saturation, respiration gated, 24 slices with two adjacent slices per packet, TR = 5.0 ms, TE = 1.4 ms, FA = 20°, NEX = 4, FOV = 64 mm x 64 mm, matrix size = 128 x 128, total acquisition time = ∼38 s) obtained immediately before or after dynamic acquisitions. Pearson correlation was calculated to statistically evaluate the agreement between these volumetric measures (a significance level of alpha = 0.05).

## Results

### Respiratory Gating and Motion Artifact Reduction

Respiratory gating significantly reduced motion artifacts associated with respiration and ensured stable, repeated acquisitions at consistent respiratory phases. Representative images (four slices, fully sampled, TR=10 ms, TE=1.4 ms, FA=25°) acquired with respiratory gating demonstrated clearer delineation of gastric geometry, particularly near the diaphragm and pylorus, compared to images acquired without gating (Fig. 5). Although sampling intervals were constrained by the duration of the respiratory cycle (typically 1–2 s per cycle), the resulting sampling rate (0.5–1 Hz) remained sufficient to capture gastric motility dynamics (∼5 cpm).

**Figure 5.**
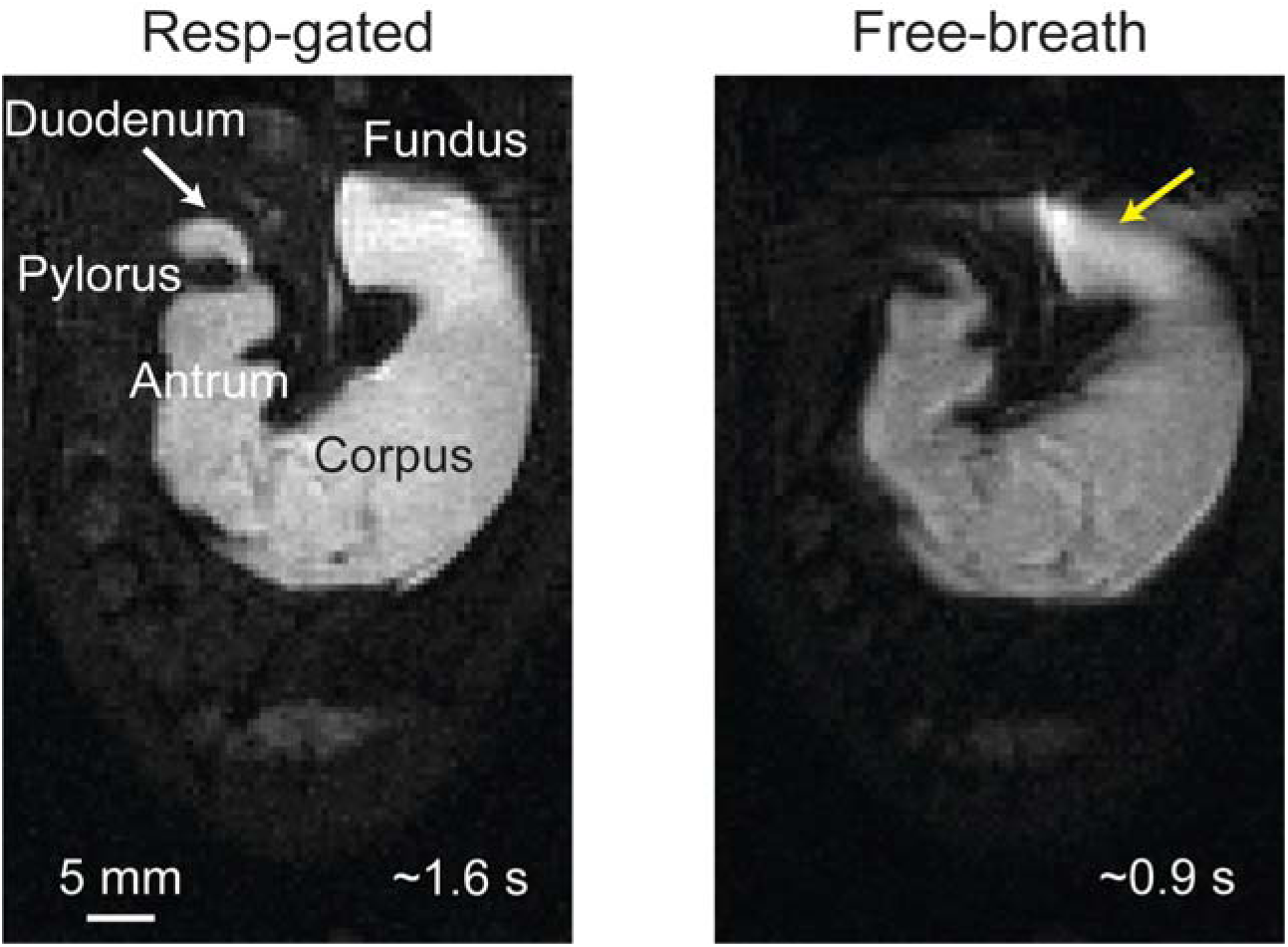
Effects of respiration gating. Fully sampled scans with (left) or without respiratory gating (right). The yellow arrow points to the fundus, where respiratory movements cause distortion. The white arrow points to the duodenum. The representative Images acquired with respiratory gating took ∼1.6 s per volume, whereas images acquired without respiratory gating took ∼0.9 s per volume.

### Effects of Spatially Non-Selective Saturation Pulses

Spatially non-selective saturation pulses enhanced gastric image quality by suppressing signals from surrounding tissues and flowing blood. Comparative images acquired with and without saturation pulses (Fig. 6) demonstrated improved intraluminal contrast due to the rapid T_1_ recovery of Gd-doped gastric contents (∼17 ms) compared to the much slower T1 recovery of surrounding tissues (> 800 ms). Importantly, saturation pulses had a negligible impact on the Gd-doped test meal (Fig. 2), and similarly for the intraluminal signal. Fig. 6 also highlights how saturation pulses effectively reduced bright blood signals and peripheral aliasing artifacts, particularly near major arteries and tissue boundaries. These improvements substantially increased the clarity of gastric structures, simplifying image segmentation and enhancing interpretability.

**Figure 6.**
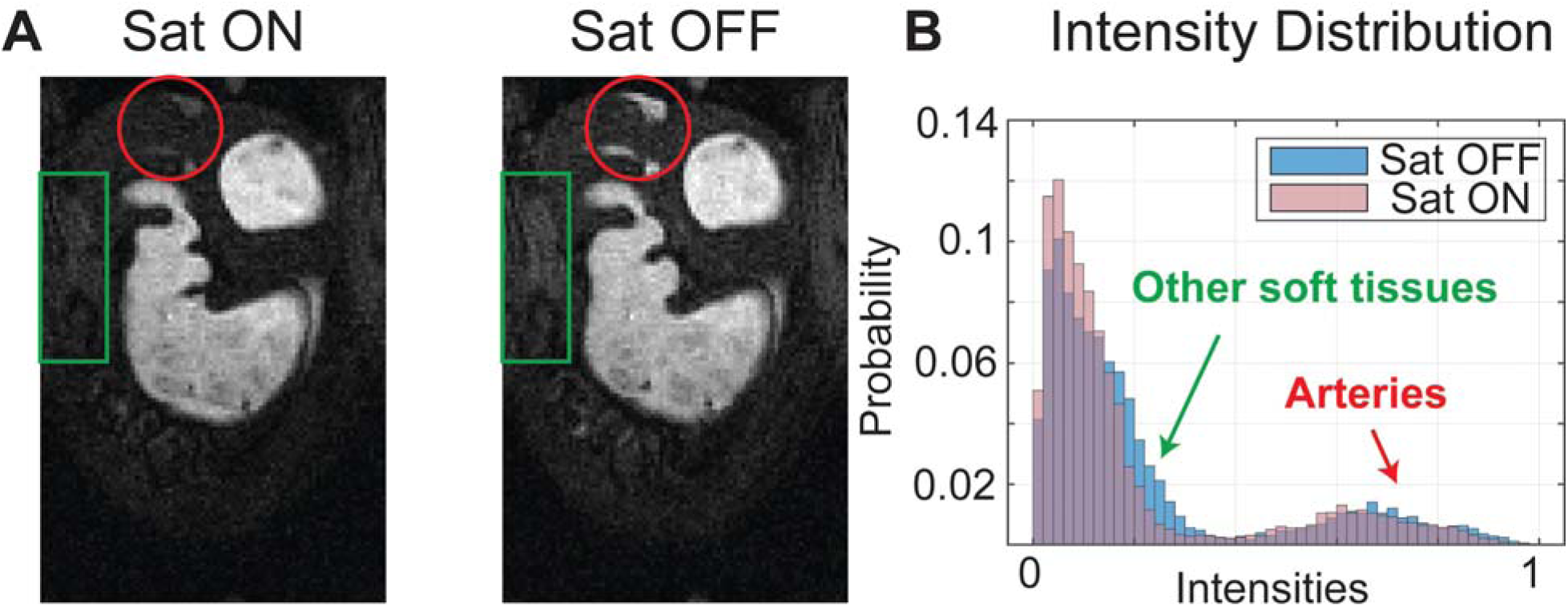
Effects of spatially non-selective saturation. **(A)** Fully sampled images with (left) or without spatially non-selective saturation pulses (right). **(B)** The distribution of normalized pixel intensities ([0 1]) from both images. Green arrows and boxes highlight other soft tissues and aliasing artifacts. Red arrows and boxes highlight arteries.

### Retrospective Reconstruction

Retrospective reconstructions from simulated undersampling scenarios (reduction factors R = 2 through 11; ACS = 12 and 24) confirmed the effectiveness and robustness of convolution-kernel-based k-t interpolation (Fig. 4). Minimal degradation in image quality was found at moderate undersampling (R = 2, 3). At higher undersampling (R = 11), noticeable increases in image blurring and reconstruction artifacts were observed (Fig. 7 and supplementary table 1). Quantitative evaluations demonstrated good reconstruction fidelity, with RE remaining low even at higher reduction factors. SSIM and PSNR metrics indicated similar trends (Fig. 7 and supplementary table 1). Gastric motility dynamics were successfully captured from retrospective reconstructions. Quantitative gastric motility metrics, such as contraction frequency, remained robustly measurable across the undersampling scenarios tested (Fig. 8 and supplementary table 2), while contraction amplitude and velocity were only mildly affected despite the statistically significant differences (Fig. 8 and supplementary table 2).

**Figure 7.**
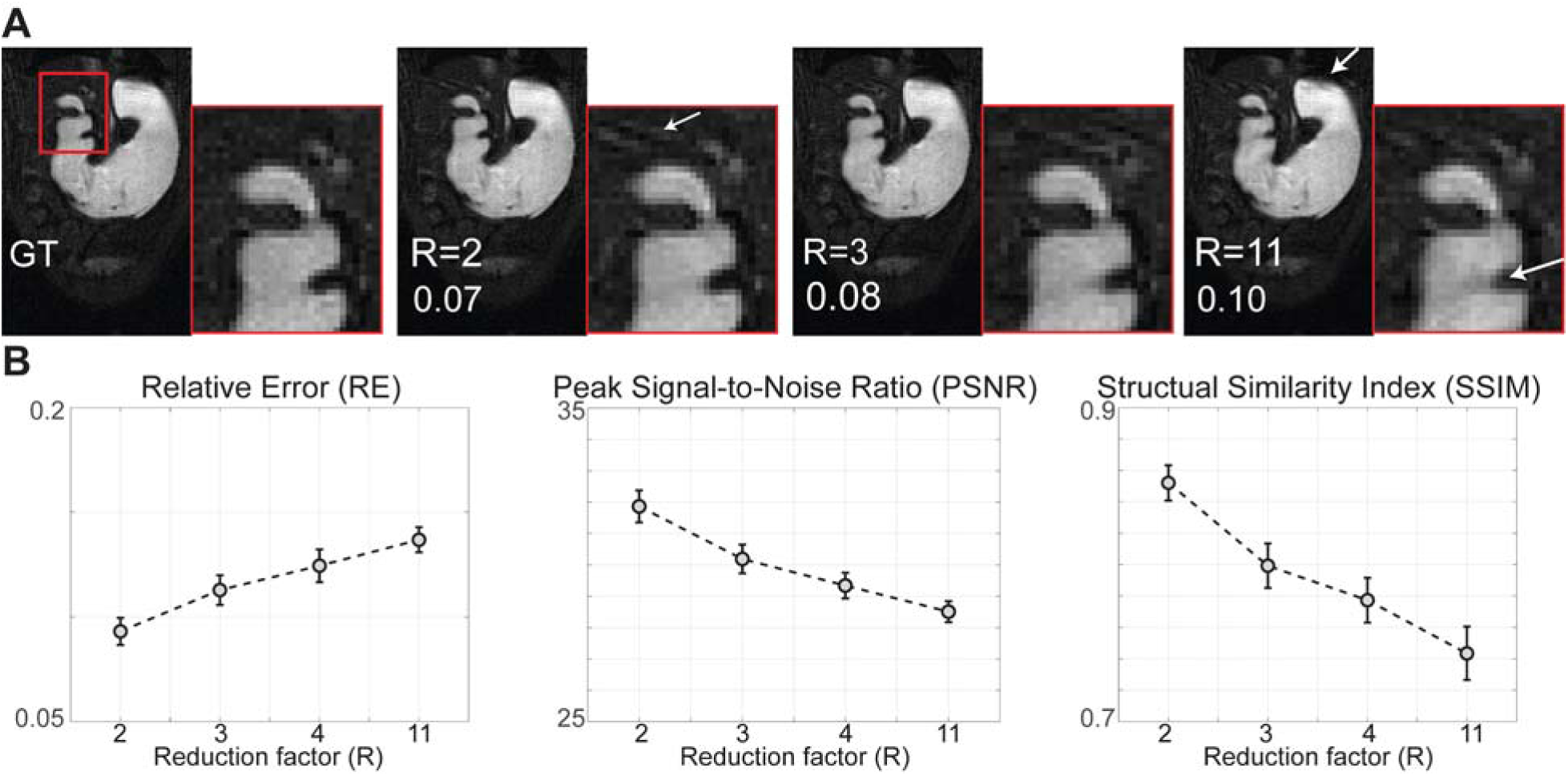
Effects of the reduction factor (R) in retrospective reconstruction. **(A)** Ground truth (GT) reference image reconstructed with fully sampled k-space, and the images reconstructed from retrospectively down-sampled k-space with R = 2, R = 3, and R = 11 (ACS = 12). Numbers on images are the relative errors (RE) between each reconstructed image and the GT image. The distal antrum, pylorus, and duodenum, highlighted by the red box, are zoomed in further. White arrows point to the artifacts. **(B)** The RE, peak signal-to-noise ratio (PSNR), and structural similarity index (SSIM) between GT and each reconstructed image with different R. The RE is calculated by the L2 norm of the intensity differences between two vectorized images divided by the L2 norm of the reference.

**Figure 8.**
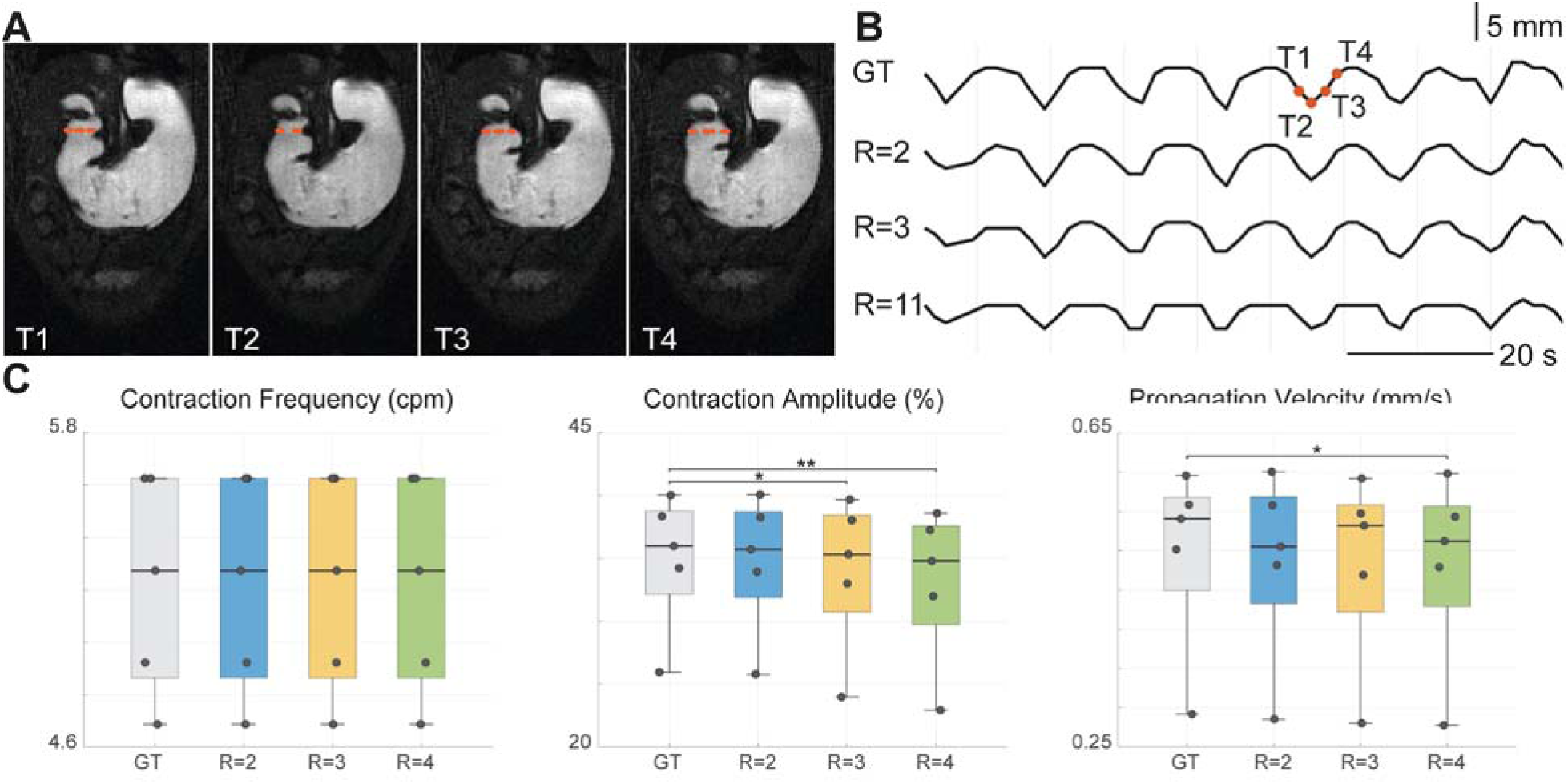
Effects of the reduction factor (R) in antral contraction quantification. **(A)** A time series of fully sampled dynamic MR images, as the ground truth (GT) reference images (T1-T4). **(B)** A time series of cross-section areas, measured from GT images and images reconstructed from retrospectively down-sampled k-space with R = 2, R = 3, and R = 11 and ACS = 12. Red dashed lines in (A) show the cross-section plotted in (B). **(C)** Quantified contraction frequency, amplitude, and propagation velocity from GT, and images reconstructed from retrospectively down-sampled k-space with R = 2, R = 3, and R = 4 (ACS = 12). Asterisk signs indicate statistical significance. * p < 0.05, ** p < 0.01.

### Accelerated Acquisition and Reconstruction

Prospective acquisitions using the proposed accelerated imaging protocol successfully achieved comprehensive stomach coverage, capturing dynamic gastric motility at high spatial and temporal resolutions. Each dataset was acquired rapidly (<3 s per image volume or 24 slices), clearly delineating GI anatomical regions such as the fundus, corpus, antrum, pylorus, and duodenum (Fig. 9A). The slice grouping scheme effectively balanced signal intensity and motion sensitivity. Compared to standard multi-slice gradient echo sequences without slice grouping, images with the slice grouping scheme were significantly less susceptible to motion artifacts, especially at high respiratory rates (e.g., > 45 cpm). A representative example (Fig. 10) illustrates that acquiring slices in packets results in a short time window (∼256 ms) for acquiring each packet and eliminates respiratory motion artifacts. Conversely, standard acquisitions without slice grouping used a longer time window (∼1,440 ms) to fill in k-space, coinciding with respiratory cycles and degrading image quality. Additionally, given a longer TR at the same FA, images acquired without slice grouping also show poorer contrast with brighter structures surrounding the GI tract. Maximum intensity projections (Fig. 9B) and the reconstructed 3D surface model (Fig. 9C) further demonstrated the high fidelity and detailed visualization of gastric morphology achieved by the proposed accelerated imaging protocol.

**Figure 9.**
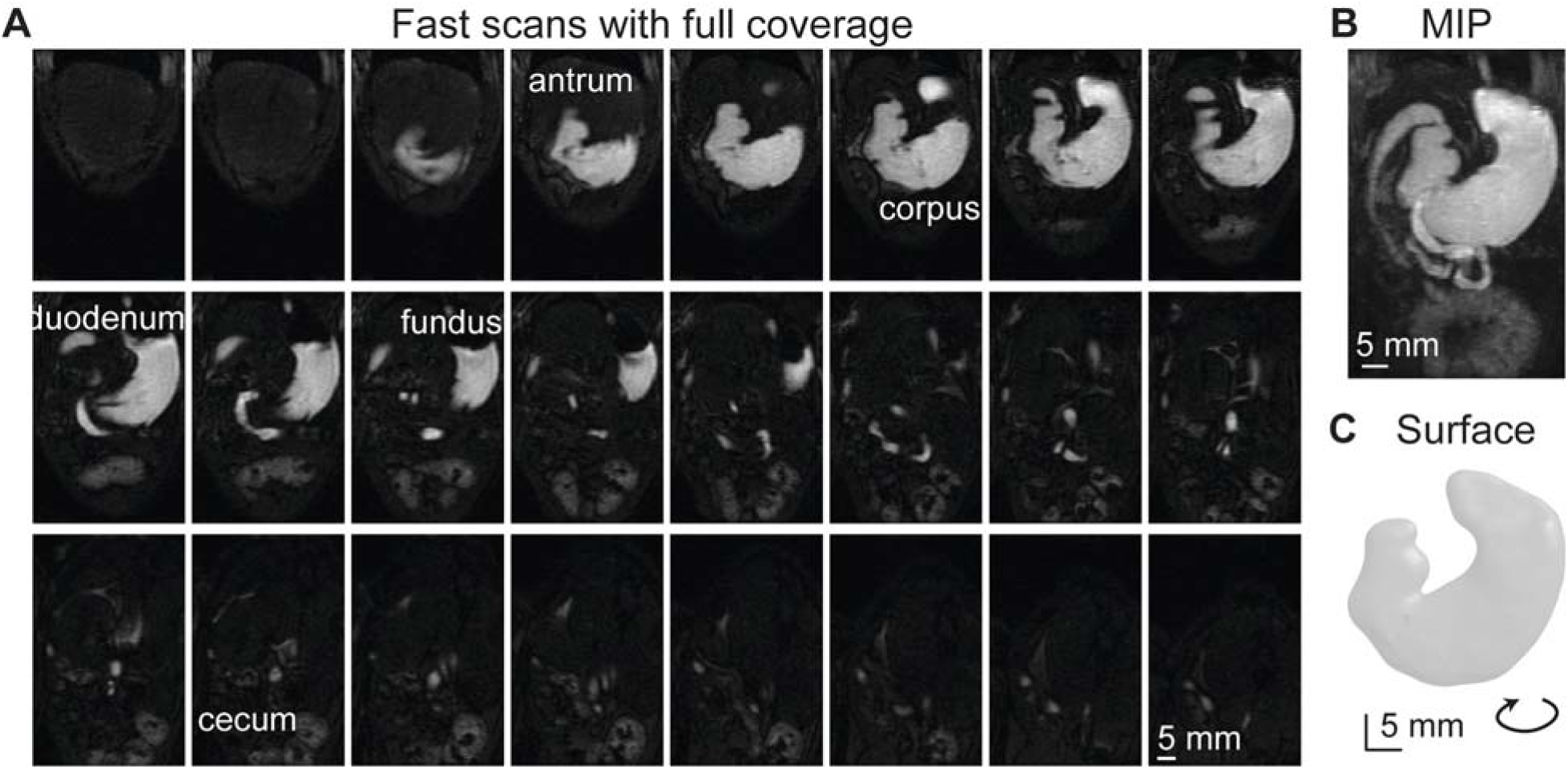
Images reconstructed from the accelerated, dynamic, under-sampled k-space cover the entire stomach and small intestines. **(A)** All 24 slices were acquired over 1.5 s, approximately one respiration cycle. **(B)** The maximum intensity projection (MIP) across 24 slices. **(C)** The corresponding 3D stomach surface model. The surface model is rotated slightly for a better visualization.

**Figure 10.**
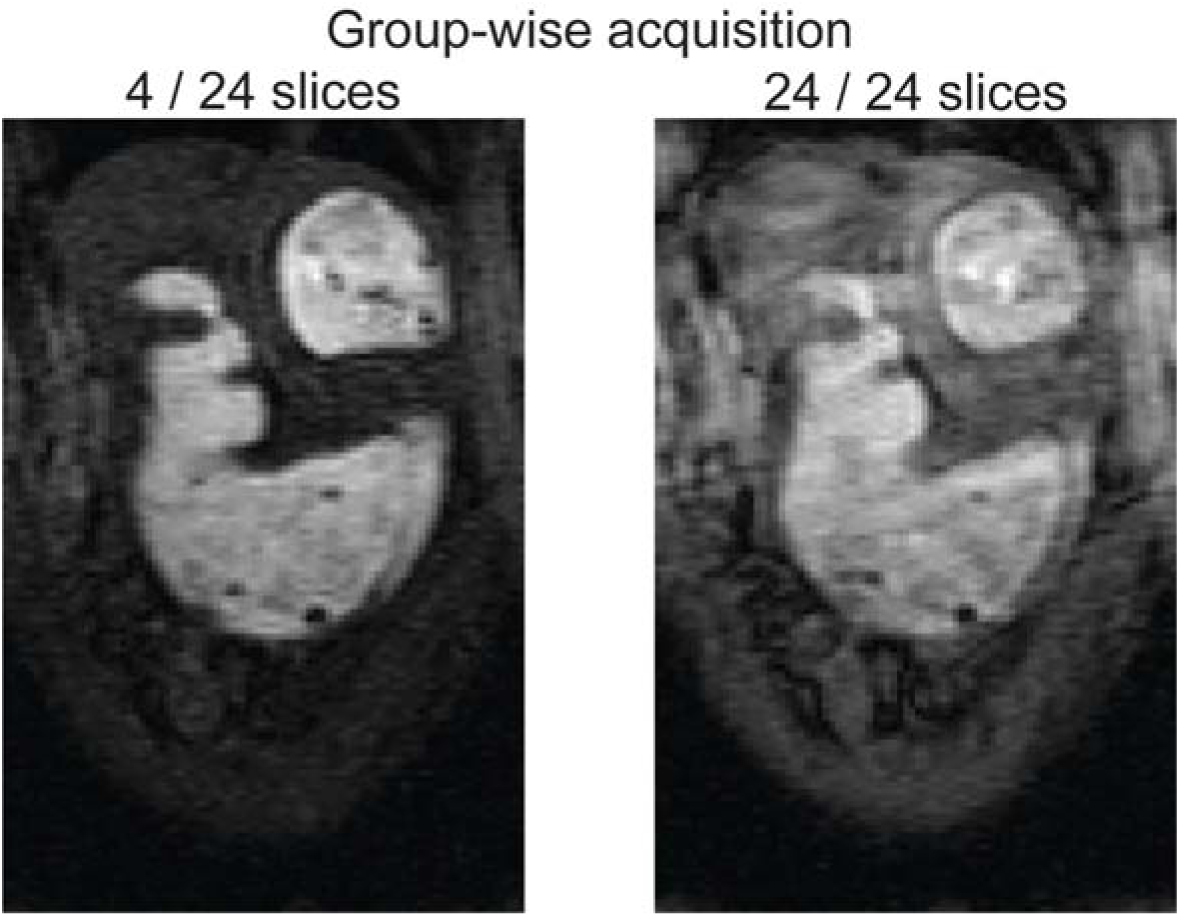
Effects of slice grouping. Left: A slice acquired within a packet of 4 adjacent slices, having an acquisition time of 256 ms per packet. Right: A comparable slice acquired within a group of 24 slices, with an acquisition time of 1440 ms. A shorter acquisition time, compared to a respiration cycle of 1-2 s, reduces motion artifacts. These representative images were acquired without respiration gating.

Prospective reconstructions accurately depict gastric dynamics and anatomy. Gastric contractions and their propagation are clearly visible along both the lesser and greater curvatures (Fig. 8). Also, see dynamic stomach motions in Supplementary Videos 1 and 2. For the group of anesthetized rats that underwent voluntary ingestion of Gd-doped test meal (n = 12), the stomach contractions have an amplitude of 21.7%+5.0%, a frequency of 5.4+0.4□cpm, and a speed of 0.56+0.11□mm/s (38), comparable to the normal ranges reported in a different group of rats (11). Gastric volume measurements (n = 12) derived from dynamic acquisitions exhibited strong agreement (r = 0.98, p < 0.001) with reference anatomical scans. These comprehensive visualizations and quantifications confirm the effectiveness of the proposed dynamic MRI approach for preclinical gastric motility studies.

## Discussion

In this paper, we describe an optimized dynamic MRI protocol and demonstrate its effectiveness for capturing whole-stomach motility in anesthetized rats. Although our method is an adaptation of existing methods rather than a radical innovation, its systematic integration of Gd-labeled meal, respiratory gating, slice grouping, and k-t undersampling and interpolation addresses several practical hurdles commonly encountered in preclinical GI MRI. Our design elements, evaluated individually and collectively, substantially improve signal uniformity, reduce motion artifacts, and yield robust measurements of gastric volumes, contraction amplitude, frequency, and velocity. The high-contrast images allow for clear delineation and easy segmentation of gastric contents, while the short scanning time (<3 s for the entire stomach) preserves sufficient temporal resolution to track gastric contractions and coordinations.

As a key advantage, this protocol achieves large spatial coverage and high temporal resolution, while maintaining the high spatial resolution necessary to resolve fine structures, such as the pylorus. In order to achieve high scanning speed, previous rat MRI studies only capture a part of the stomach (11), underestimating total gastric volume and lacking the ability to evaluate motility across the entire stomach. Here, full-stomach coverage plus submillimeter resolution and <3 s temporal resolution promises to accurately track gastric wall motion and uncover regional differences in contractile activity. Saturation pulses selectively suppress extraluminal signals, exploiting the significant T_1_ difference between Gd-doped contents (∼17 ms) and tissues (>800 ms). In addition, respiratory gating aligns acquisition, minimizing image blurring and motion artifacts. The resultant dataset can report anatomical, volumetric, and motility metrics at global, regional, and subregional levels, enabling detailed evaluation and mapping of gastric motor function (21,39).

Our study extends the practical applicability of k-t interpolation. Single-coil GI MRI is challenging because typical parallel imaging frameworks require multiple coils bearing different spatial sensitivities. However, because the Gd-doped meal appears relatively homogeneous, an effective local “sensitivity” pattern emerges in the lumen, supporting the assumption of a convolutional kernel shift-invariant in k space. This strategy allows for accelerated data acquisition while retaining motion details. Although this method is similar to k-t GRAPPA initially introduced for dynamic cardiac imaging (41), its application in small-animal GI MRI is our unique contribution, adding a compelling dimension to enhance the practicality of preclinical imaging of gastric motility.

By offering a noninvasive readout of gastric motility patterns, our protocol has broad implications. It can be used to test new prokinetic agents or neuromodulatory drugs in rodent models (44,45), where effects can be monitored in real time. Researchers exploring vagus nerve stimulation, optogenetic or chemogenetic stimulation, or target lesioning can likewise quantify how neural control or ablation affects gastric motility. Our method can be used in disease models to interrogate pathophysiology in GI disorders, such as gastroparesis, and provide mechanistic insights into how various gastric regions and their motor events contribute to delayed gastric emptying. Our method can also provide a deeper understanding of gastric physiology. By varying the composition of ingested test meals (e.g., different nutrient profiles or meal volume) or manipulating gastric pH and secretions, their effects become quantifiable through the volumetric and dynamic measures obtained with gastric MRI. Due to the non-invasive nature of MRI, it is plausible to collect repeated, longitudinal data in the same animal, benefiting chronic studies and supporting robust experimental designs as well as better translation to humans.

Applying advanced coil designs, parallel imaging, or compressed sensing may further shorten acquisition time or enhance spatial resolution, aiding studies targeting faster dynamics or finer structures. Comparing gastric volume or motility measurements derived from the proposed GI MRI protocol with other terminal or invasive reference methods (6–9) in rodents would further validate its accuracy and robustness. Extending the method in rodent disease models, such as chemically induced gastroparesis, would demonstrate its utility for diagnosing and monitoring clinically relevant phenotypes. Each of these directions holds potential to refine the protocol and expand its scope, propelling MRI-based motility assessment toward an indispensable standard in preclinical gastroenterology.

This study has limitations. The use of anesthesia remains a challenge. Even mild anesthetics can reduce gastric contractility (9,38), and different agents or doses may affect motility patterns differently (38,46). Translating the MRI protocol from anesthetized conditions to awake conditions requires caution and fine-tuning (38). Another limitation is that mixing gastric juice with Gd-labeled gel in vivo can complicate the differentiation between meal volume and secretions, potentially confounding exact measures of emptying. Further innovations could incorporate multi-contrast approaches or T_1_/T_2_ mapping to parse out the role of secretions (22,23), differentiating labeled meal content from unlabeled fluid in the lumen. Lastly, while we focus on dynamic movements of the stomach, intragastric pressure is not addressed in our current protocol, but worth investigating.

## Conclusion

The protocol described here provides an accessible and standardized framework for real-time MRI of rat gastric motility. Its combination of respiratory gating, saturation pulses, accelerated imaging, and semi-solid contrast meal preparation overcomes many prior constraints, facilitating high-fidelity visualization of gastric motor function. From pharmacological testing to disease characterization, the method paves the way for comprehensive, repeated, and quantitative studies of rodent gastric physiology that can inform and complement human research.

## Abbreviations

ACS: Auto-calibration signal
cpm: Cycles per minute
FOV: Field of view
FA: Flip angle
GI: Gastrointestinal
PSNR: Peak signal-to-noise ratio
PE: Phase-encoding
R: Reduction factor
RE: Relative error
SSIM: Structural Similarity Index

## Notes

### Competing Interest Statement

The authors have declared no competing interest.

## References

1. Greenwood-Van Meerveld B, Johnson AC, Grundy D. Gastrointestinal Physiology and Function. In: Handbook of Experimental Pharmacology. 2017. p. 1–16.

2. Goyal RK, Guo Y, Mashimo H. Advances in the physiology of gastric emptying. Neurogastroenterol Motil. 2019 Apr;31(4):e13546.

3. Camilleri M, Sanders KM. Gastroparesis. Gastroenterology. 2022 Jan;162(1):68–87.e1.

4. Camilleri M, Chedid V, Ford AC, Haruma K, Horowitz M, Jones KL, et al. Gastroparesis. Nat Rev Dis Primer. 2018 Nov 1;4(1):41.

5. Di Natale MR, Athavale ON, Wang X, Du P, Cheng LK, Liu Z, et al. Functional and anatomical gastric regions and their relations to motility control. Neurogastroenterol Motil. 2023 Sep;35(9):e14560.

6. Zheng J, Dobner A, Babygirija R, Ludwig K, Takahashi T. Effects of repeated restraint stress on gastric motility in rats. Am J Physiol-Regul Integr Comp Physiol [Internet]. 2009 May [cited 2025 Feb 25];296(5):R1358–65. Available from: https://journals.physiology.org/doi/full/10.1152/ajpregu.90928.2008

7. Torjman MC, Joseph JI, Munsick C, Morishita M, Grunwald Z. Effects of isoflurane on gastrointestinal motility after brief exposure in rats. Int J Pharm. 2005 Apr 27;294(1–2):65– 71.

8. Athavale ON, Di Natale MR, Avci R, Clark AR, Furness JB, Cheng LK, et al. Mapping the rat gastric slow-wave conduction pathway: bridging in vitro and in vivo methods, revealing a loosely coupled region in the distal stomach. Am J Physiol Gastrointest Liver Physiol. 2024 Aug 1;327(2):G254–66.

9. Tomaselli L, Sciullo M, Fulton S, Yates BJ, Fisher LE, Ventura V, et al. Isoflurane anesthesia suppresses gastric myoelectric power in the ferret. Neurogastroenterol Motil. 2024 Mar;36(3):e14749.

10. Janssen P, Nielsen MA, Hirsch I, Svensson D, Gillberg PG, Hultin L. A novel method to assess gastric accommodation and peristaltic motility in conscious rats. Scand J Gastroenterol. 2008 Jan;43(1):34–43.

11. Lu KH, Cao J, Oleson ST, Powley TL, Liu Z. Contrast-Enhanced Magnetic Resonance Imaging of Gastric Emptying and Motility in Rats. IEEE Trans Biomed Eng. 2017 Nov;64(11):2546–54.

12. Chavero-Pieres M, Viola MF, Appeltans I, Abdurahiman S, Gsell W, Matteoli G, et al. Magnetic resonance imaging as a non-invasive tool to assess gastric emptying in mice. Neurogastroenterol Motil. 2023 Feb;35(2):e14490.

13. Marciani L. Assessment of gastrointestinal motor functions by MRI: a comprehensive review. Neurogastroenterol Motil. 2011 May;23(5):399–407.

14. Spiller R, Marciani L. Intraluminal Impact of Food: New Insights from MRI. Nutrients. 2019 May 23;11(5):1147.

15. Coleman NS, Marciani L, Blackshaw E, Wright J, Parker M, Yano T, et al. Effect of a novel 5-HT3 receptor agonist MKC-733 on upper gastrointestinal motility in humans. Aliment Pharmacol Ther. 2003 Nov 15;18(10):1039–48.

16. de Zwart IM, Haans JJL, Verbeek P, Eilers PHC, de Roos A, Masclee AAM. Gastric accommodation and motility are influenced by the barostat device: Assessment with magnetic resonance imaging. Am J Physiol Gastrointest Liver Physiol. 2007 Jan;292(1):G208–214.

17. Haans JJL, de Zwart IM, Eilers PHC, Reiber JHC, Doornbos J, de Roos A, et al. Gastric volume changes in response to a meal: validation of magnetic resonance imaging versus the barostat. J Magn Reson Imaging JMRI. 2011 Sep;34(3):685–90.

18. Lu KH, Liu Z, Jaffey D, Wo JM, Mosier KM, Cao J, et al. Automatic assessment of human gastric motility and emptying from dynamic 3D magnetic resonance imaging. Neurogastroenterol Motil. 2022 Jan;34(1):e14239.

19. Sclocco R, Nguyen C, Staley R, Fisher H, Mendez A, Velez C, et al. Non-uniform gastric wall kinematics revealed by 4D Cine magnetic resonance imaging in humans. Neurogastroenterol Motil. 2021 Aug;33(8):e14146.

20. Menys A, Hoad C, Spiller R, Scott SM, Atkinson D, Marciani L, et al. Spatio-temporal motility MRI analysis of the stomach and colon. Neurogastroenterol Motil. 2019 May;31(5):e13557.

21. Wang X, Alkaabi F, Choi M, Di Natale MR, Scheven UM, Noll DC, et al. Surface mapping of gastric motor functions using MRI: a comparative study between humans and rats. Am J Physiol-Gastrointest Liver Physiol [Internet]. 2024 Sep [cited 2024 Dec 17];327(3):G345–59. Available from: https://journals.physiology.org/doi/full/10.1152/ajpgi.00045.2024

22. Goetze O, Treier R, Fox M, Steingoetter A, Fried M, Boesiger P, et al. The effect of gastric secretion on gastric physiology and emptying in the fasted and fed state assessed by magnetic resonance imaging. Neurogastroenterol Motil. 2009 Jul;21(7):725–e42.

23. Sauter M, Curcic J, Menne D, Goetze O, Fried M, Schwizer W, et al. Measuring the interaction of meal and gastric secretion: a combined quantitative magnetic resonance imaging and pharmacokinetic modeling approach. Neurogastroenterol Motil. 2012 Jul;24(7):632–8, e272-273.

24. Khalaf A, Hoad CL, Menys A, Nowak A, Taylor SA, Paparo S, et al. MRI assessment of the postprandial gastrointestinal motility and peptide response in healthy humans. Neurogastroenterol Motil. 2018 Jan;30(1).

25. Menys A, Butt S, Emmanuel A, Plumb AA, Fikree A, Knowles C, et al. Comparative quantitative assessment of global small bowel motility using magnetic resonance imaging in chronic intestinal pseudo-obstruction and healthy controls. Neurogastroenterol Motil. 2016 Mar;28(3):376–83.

26. Lu KH, Cao J, Oleson S, Ward MP, Phillips RJ, Powley TL, et al. Vagus nerve stimulation promotes gastric emptying by increasing pyloric opening measured with magnetic resonance imaging. Neurogastroenterol Motil. 2018 Oct;30(10):e13380.

27. Lu KH, Cao J, Phillips R, Powley TL, Liu Z. Acute effects of vagus nerve stimulation parameters on gastric motility assessed with magnetic resonance imaging. Neurogastroenterol Motil. 2020 Jul;32(7):e13853.

28. Griswold MA, Jakob PM, Heidemann RM, Nittka M, Jellus V, Wang J, et al. Generalized autocalibrating partially parallel acquisitions (GRAPPA). Magn Reson Med. 2002 Jun;47(6):1202–10.

29. Pruessmann KP, Weiger M, Scheidegger MB, Boesiger P. SENSE: sensitivity encoding for fast MRI. Magn Reson Med. 1999 Nov;42(5):952–62.

30. Uecker M, Lai P, Murphy MJ, Virtue P, Elad M, Pauly JM, et al. ESPIRiT--an eigenvalue approach to autocalibrating parallel MRI: where SENSE meets GRAPPA. Magn Reson Med. 2014 Mar;71(3):990–1001.

31. Lin CY, Fessler JA. Efficient Dynamic Parallel MRI Reconstruction for the Low-Rank Plus Sparse Model. IEEE Trans Comput Imaging. 2019 Mar;5(1):17–26.

32. Majumdar A, Ward RK, Aboulnasr T. Compressed sensing based real-time dynamic MRI reconstruction. IEEE Trans Med Imaging. 2012 Dec;31(12):2253–66.

33. Caballero J, Price AN, Rueckert D, Hajnal JV. Dictionary learning and time sparsity for dynamic MR data reconstruction. IEEE Trans Med Imaging. 2014 Apr;33(4):979–94.

34. Otazo R, Candès E, Sodickson DK. Low-rank plus sparse matrix decomposition for accelerated dynamic MRI with separation of background and dynamic components. Magn Reson Med. 2015 Mar;73(3):1125–36.

35. Yaman B, Weingärtner S, Kargas N, Sidiropoulos ND, Akçakaya M. Locally Low-Rank Tensor Regularization for High-Resolution Quantitative Dynamic MRI. Int Workshop Comput Adv Multi-Sens Adapt Process Int Workshop Comput Adv Multi-Sens Adapt Process. 2017 Dec;2017.

36. Pal A, Rathi Y. A review and experimental evaluation of deep learning methods for MRI reconstruction. J Mach Learn Biomed Imaging. 2022 Mar;1:001.

37. Oscanoa JA, Middione MJ, Alkan C, Yurt M, Loecher M, Vasanawala SS, et al. Deep Learning-Based Reconstruction for Cardiac MRI: A Review. Bioeng Basel Switz. 2023 Mar 6;10(3):334.

38. Wang X, Alkaabi F, Cornett A, Choi M, Scheven UM, Di Natale MR, et al. Magnetic Resonance Imaging of Gastric Motility in Conscious Rats. Neurogastroenterol Motil. 2024 Dec 31;e14982.

39. Wang X, Cao J, Han K, Choi M, She Y, Scheven UM, et al. Diffeomorphic Surface Modeling for MRI-Based Characterization of Gastric Anatomy and Motility. IEEE Trans Biomed Eng. 2023 Jul;70(7):2046–57.

40. Haldar JP, Setsompop K. Linear Predictability in MRI Reconstruction: Leveraging Shift-Invariant Fourier Structure for Faster and Better Imaging. IEEE Signal Process Mag [Internet]. 2020 Jan [cited 2025 Mar 31];37(1):69–82. Available from: https://www.ncbi.nlm.nih.gov/pmc/articles/PMC7971148/

41. Huang F, Akao J, Vijayakumar S, Duensing GR, Limkeman M. k-t GRAPPA: a k-space implementation for dynamic MRI with high reduction factor. Magn Reson Med. 2005 Nov;54(5):1172–84.

42. Qu X, Guo D, Ning B, Hou Y, Lin Y, Cai S, et al. Undersampled MRI reconstruction with patch-based directional wavelets. Magn Reson Imaging [Internet]. 2012 Sep 1 [cited 2025 Feb 28];30(7):964–77. Available from: https://www.sciencedirect.com/science/article/pii/S0730725X12000604

43. Wang Z, Bovik AC, Sheikh HR, Simoncelli EP. Image quality assessment: from error visibility to structural similarity. IEEE Trans Image Process [Internet]. 2004 Apr [cited 2025 Mar 1];13(4):600–12. Available from: https://ieeexplore.ieee.org/document/1284395

44. Viramontes BE, Kim DY, Camilleri M, Lee JS, Stephens D, Burton DD, et al. Validation of a stable isotope gastric emptying test for normal, accelerated or delayed gastric emptying. Neurogastroenterol Motil [Internet]. 2001 [cited 2023 Nov 21];13(6):567–74. Available from: https://onlinelibrary.wiley.com/doi/abs/10.1046/j.1365-2982.2001.00288.x

45. Parthasarathy G, Ravi K, Camilleri M, Andrews C, Szarka LA, Low PA, et al. Effect of neostigmine on gastroduodenal motility in patients with suspected gastrointestinal motility disorders. Neurogastroenterol Motil. 2015 Dec;27(12):1736–46.

46. Qualls-creekmore E, Tong M, Holmes GM. Gastric emptying of enterally administered liquid meal in conscious rats and during sustained anaesthesia. Neurogastroenterol Motil [Internet]. 2010 [cited 2025 Feb 25];22(2):181–5. Available from: https://onlinelibrary.wiley.com/doi/abs/10.1111/j.1365-2982.2009.01393.x

